# Self-limiting population suppression gene drive in the West Nile vector mosquito, *Culex quinquefasciatus*

**DOI:** 10.1101/2025.07.16.665070

**Authors:** Xuechun Feng, Jinying Ding, Yiran Liu, Víctor López Del Amo, Valentino M. Gantz, Xuexin Chen, Jackson Champer, Feng Liu

## Abstract

*Culex* mosquitoes transmit major pathogens including West Nile virus, encephalitis, filariasis, and avian malaria, threatening public health, poultry, and ecosystems. We engineered a CRISPR-based population suppression gene drive targeting a conserved exon of the *doublesex* (*dsx*) gene. The drive incorporates a recoded *dsxM* segment to preserve male function while converting genetic females into sterile intersexes, enabling male-biased propagation and removal of fertile females. It achieves super-Mendelian inheritance (∼71%) and generates partially dominant sterile resistance alleles via end-joining, resulting in intersex phenotypes with reduced fertility and hatchability. Modeling predicts that this RIDD (Release of Insects carrying a Dominant-sterile Drive) system can suppress populations at low intrinsic growth rates and release ratios, outperforming SIT and fs-RIDL strategies in persistence and efficiency, with further gains achievable through improved cleavage rates. This study establishes a self-limiting gene drive framework for *Culex* suppression, highlightling the potential of targeting conserved sex-determination pathways for sustainable vector control.

## INTRODUCTION

Homing gene drives are genetic elements capable of spreading through populations by biasing their inheritance^1–3^. Among the platforms developed, CRISPR-based gene drives are the most advanced. These systems typically consist of Cas9 and guide RNA (gRNA) components integrated at a specific genomic locus. Upon expression, the Cas9/gRNA complex induces a double-strand break at the corresponding wild-type allele, which is subsequently repaired via homology-directed repair (HDR), copying the drive allele into the break site. This process converts heterozygotes into homozygotes in the germline, resulting in super-Mendelian (>50%) inheritance in subsequent generations^4^. CRISPR-based gene drives have been successfully demonstrated across diverse organisms, including flies^4–7^, bacteria^8^, yeast^8,9^, mice^10^ and mosquitoes from *Anopheline^11–16^*, *Aedes^17^*, and *Culex^18^* genera.

In mosquitoes, gene drives have been engineered for two primary purposes: population modification, in which transgenic mosquitoes carrying anti-pathogen effectors replace wild-type populations^14,19^, and population suppression, which disrupts genes essential for reproduction or sex determination^12,14^. While gene drives hold transformative promise for controlling vector-borne diseases, their capacity for uncontrolled spread, even from low initial introduction frequency, has raised significant ecological, ethical, and regulatory concerns^20–23^. In particular, the potential for transboundary dispersal and unintended gene flow remains a major challenge for safe deployment ^24,25^.

To mitigate these concerns, self-limiting gene drives have been developed. These systems spread transiently due to high release thresholds or intrinsic fitness costs, limiting their long-term persistence and making them more suitable for confined field trials^26–28^. One such system is RIDD (Release of Insects carrying a Dominant-sterile Drive), an improved self-limiting population suppression strategy that integrates concepts from fs-RIDL (Release of Insects carrying a Dominant Lethal) with CRISPR-based drives. In RIDD, drive-carrying males bias inheritance of the construct, while female progeny inheriting either the drive or nonfunctional resistance alleles (generated via end-joining) are rendered sterile or non-viable^6,29^. A proof-of-concept RIDD system was demonstrated in *Drosophila melanogaster*, where the drive targeted the female-specific exon of the *doublesex* (*dsx*) gene, a conserved regulator of insect sexual development, causing dominant female sterility and intersex phenotypes, and repeated release of RIDD males in laboratory cages led to progressive suppression and eventual population collapse^6^. This system is potentially substantially more powerful than non-drive systems such as SIT (Sterile Insect Technique) and fs-RIDL.

As a terminal effector of the sex determination cascade, alternative splicing of *dsx* directs male and female differentiation, making it a compelling target for suppression drives^31–36^. Although disruption early in the female-specific exon of *dsx* induces dominant female sterility in *Drosophila^6,30^*, studies in *Anopheles* mosquitoes suggest that insertions at this site are recessive sterile^12,14^. In *An. gambiae*, a CRISPR-based gene drive targeting the female-specific exon of *dsx* achieved >90% inheritance and drove population collapse in cage trials^14^. Similarly, in *An. stephensi*, targeting the same exon yielded ∼75% drive efficiency, which increased to ∼100% in a *vasa*-Cas9 background^12^. These results underscore the value of *dsx* as a target for gene drive-based suppression. Nonetheless, drive performance can be compromised by maternal Cas9 deposition and resistance allele formation via end-joining, which may be mitigated by optimizing germline-specific expressions and multiplexed gRNA designs^1,7,37–39^.

In *Culex* mosquitoes, *dsx* function has been preliminarily investigated^40^, but the effect of transgene insertions (dominant or recessive effect) remains unclear. Moreover, the primary sex-determination signal and upstream regulators controlling *dsx* splicing in *Culex* are still poorly understood^41,42^. Although one study in *Culex* demonstrated proof-of-concept gene drives targeting the *white* and *kmo* loci^43^, a functional suppression drive has not yet been established. *Culex quinquefasciatus*, in particular, is a major vector of West Nile virus, lymphatic filariasis, and avian malaria^44,45^, spreading pathogens affecting humans, livestock, companion animals, and endangered wildlife^46,47^, with increasing relevance for island conservation. Compounding the challenge is the rise of insecticide resistance in *Culex* populations, which undermines the efficacy of conventional control strategies^48,49^.

In this study, we developed a CRISPR-based, self-limiting suppression gene drive targeting the *dsx* locus in *Cx. quinquefasciatus*. Our construct disrupts a conserved exon (exon 4) shared by male and female isoforms and incorporates a recoded *dsxM* fragment to preserve normal male function, enabling males to remain fertile and propagate the drive. To enhance efficiency, we employed a *nanos* germline promoter and multiplexed gRNA design. This system enables drive-carrying males to spread the construct while progressively reducing the number of fertile females through the generation of sterile drive-carrying intersexes and partially sterile non-drive intersexes. These findings indicate that *dsx* disruption induces dominant sterility in *Culex*, mirroring observations in the *Drosophila* RIDD system^6^. Thus, we designated this system as a RIDD system for *Culex*. Despite moderate inheritance efficiency, modeling shows that the drive can suppress populations under low intrinsic growth rates and release ratios, outperforming SIT and fs-RIDL strategies, though higher cleavage rates are needed before more robust populations can be targeted. Importantly, the fitness cost associated with intersex phenotypes facilitates natural drive elimination, reinforcing its self-limiting behavior and improving biosafety. This work establishes a fundamental, scalable gene drive strategy for *Culex* suppression, with promising implications for vector control efforts.

## RESULTS

### Generation of a CRISPR-based suppression gene drive targeting the *Culex dsx* locus

We engineered a homing gene drive system in *Cx. quinquefasciatus* by targeting exon 4 of the *doublesex* (*dsx*) gene, a region shared by both male and female isoforms (**Fig.1A**). The drive construct incorporates a recoded sequence comprising partial exon 4 and a *dsxM* cDNA fragment, inserted precisely at the 3’ junction of the gRNA cleavage site, followed by a 3’UTR from *An. gambiae kynurenine hydroxylase* (*kh*) gene (*LOC1276617*). This design enables the formation of an in-frame transcript that preserves male-specific *dsxM* function while evading recognition by the Cas9/gRNA complex. The strategy aims to convert genetic females carrying the drive into sterile intersexes with masculinized traits.

**Figure 1:**
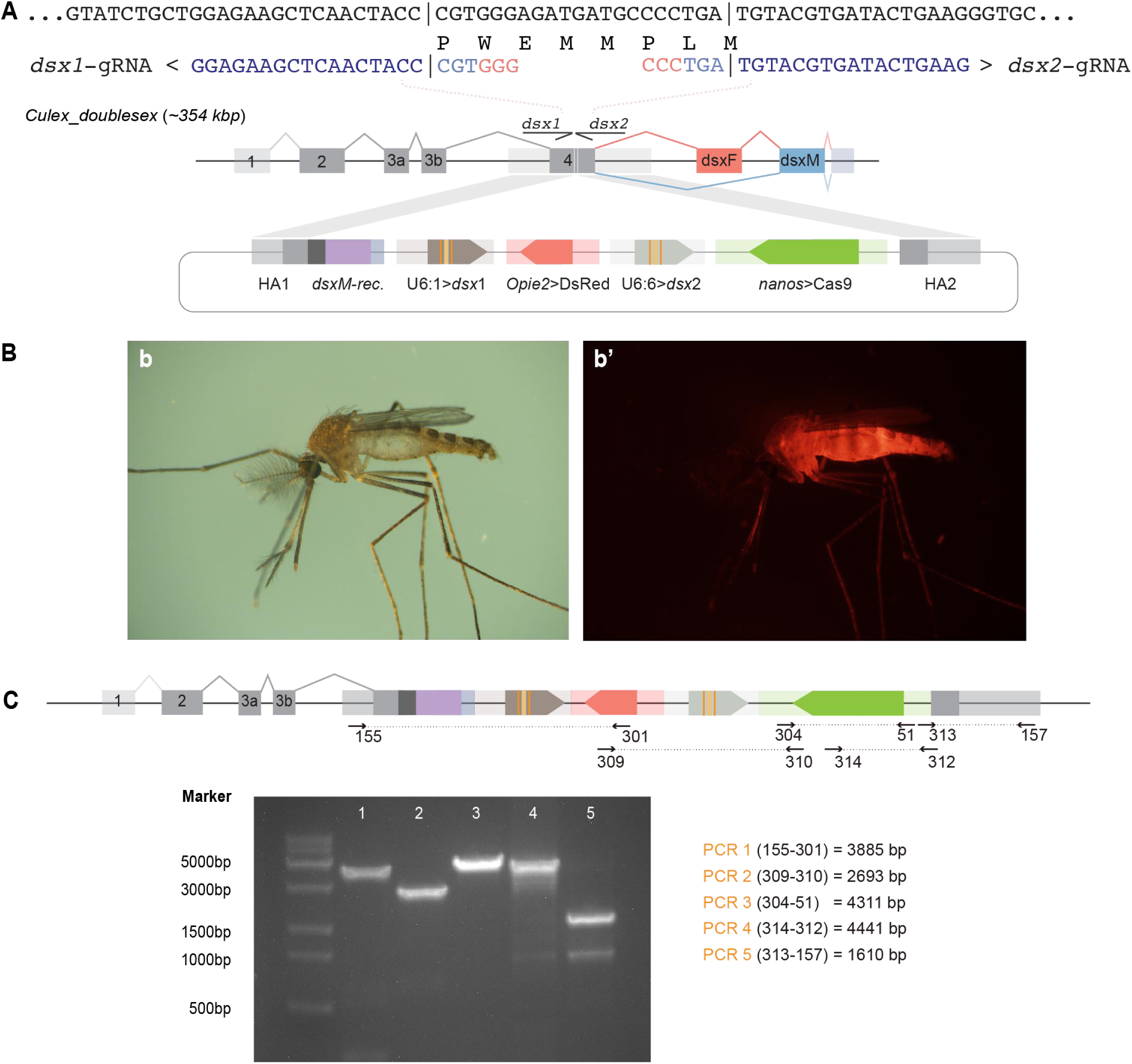
Generation and molecular validation of a homing suppression gene drive targeting the *doublesex* (*dsx*) locus in *Cx. quinquefasciatus*. A: Schematic of the *dsx* gene illustrating alternative splicing isoforms: the male-specific (*dsxM*) and female-specific (*dsxF*) transcripts. Exons shared by both isoforms are shaded in gray; the male-specific exon is labeled blue (*dsxM*), and the female-specific exons in red (*dsxF*). Exons and introns are not drawn to scale. Two guide RNAs (*dsx-1* and *dsx-2*), oriented in opposite directions, target the conserved exon with their sequences shown above. The gene drive construct includes the following elements: *nanos*-Cas9, U6:1-*dsx1* gRNA, U6:2-*dsx2* gRNA, an *Opie2*-DsRed fluorescent marker, a recoded *dsxM* cDNA fragment followed by the *An. gambiae kh* 3’UTR, and two flanking homology arms (HA1 and HA2). The recoded *dsxM* fragment is inserted immediately downstream of HA1. **B**: Representative images of transgenic *Culex* mosquitoes under brightfield and fluorescence microscopy, showing strong, ubiquitous DsRed expression. **C:** Gel electrophoresis of diagnostic PCR amplicons confirming precise insertion of the drive construct at the *dsx* locus, using specific primer sets.

To minimize the formation of resistance alleles, Cas9 expression was driven by the germline-specific *nanos* promoter. Two previously validated Pol III promoters, U6:1 and U6:6, were used to drive expression of a pair of closely spaced, oppositely oriented gRNAs targeting exon 4, with a 21 bp interval between their cleavage sites. This gRNA configuration was designed to enhance cleavage efficiency and precision. The drive cassette also includes a DsRed fluorescent marker driven by the *Opie2* promoter and is flanked by ∼1.5kb homology arms corresponding to the *dsx* locus (**Fig.1A**).

To generate transgenic mosquitoes, approximately 500 *Culex* eggs were microinjected with the gene drive plasmid at a final concentration of 300 ng/ul. Surviving G0 individuals were sexed and outcrossed to wild-type mosquitoes. From ∼1000 G1 progeny derived from the male pool, six transgenic individuals were recovered based on strong, body-wide DsRed fluorescence (**Fig. 1B, Supplementary Data 1**). One male, confirmed to carry the correctly integrated construct and exhibiting a normal phenotype, was selected to establish a stable drive line by crossing with 10 wild-type females. The line was maintained through successive generations by selecting DsRed-positive males and outcrossing to wild-type females. Precise integration of the drive cassette at the *dsx* locus was confirmed by PCR and Sanger sequencing (**Fig.1C**).

### Super-Mendelian inheritance of the gene drive

To evaluate the efficiency of the homing gene drive, we initiated experiments by crossing 10 drive-carrying males with 10 wild-type females to generate G1 heterozygous carriers. Approximately 20 single-pair crosses between G1 heterozygous males and wild-type females were established to assess gene drive inheritance in the G2 generation. Two independent experimental batches conducted at different times served as biological replicates, and data were pooled for analysis. At the adult stage, G2 individuals were screened for DsRed fluorescence and morphological phenotypes indicative of various gene editing outcomes (**Fig. 2A**). DsRed-positive individuals confirmed inheritance of the gene drive construct and exhibited either male or intersex phenotypes, depending on their genetic sex. Genetic males carrying the gene drive retained normal male characteristics, while genetic females inheriting the drive with the recoded *dsxM* sequence developed masculinized intersex phenotypes.

**Figure 2:**
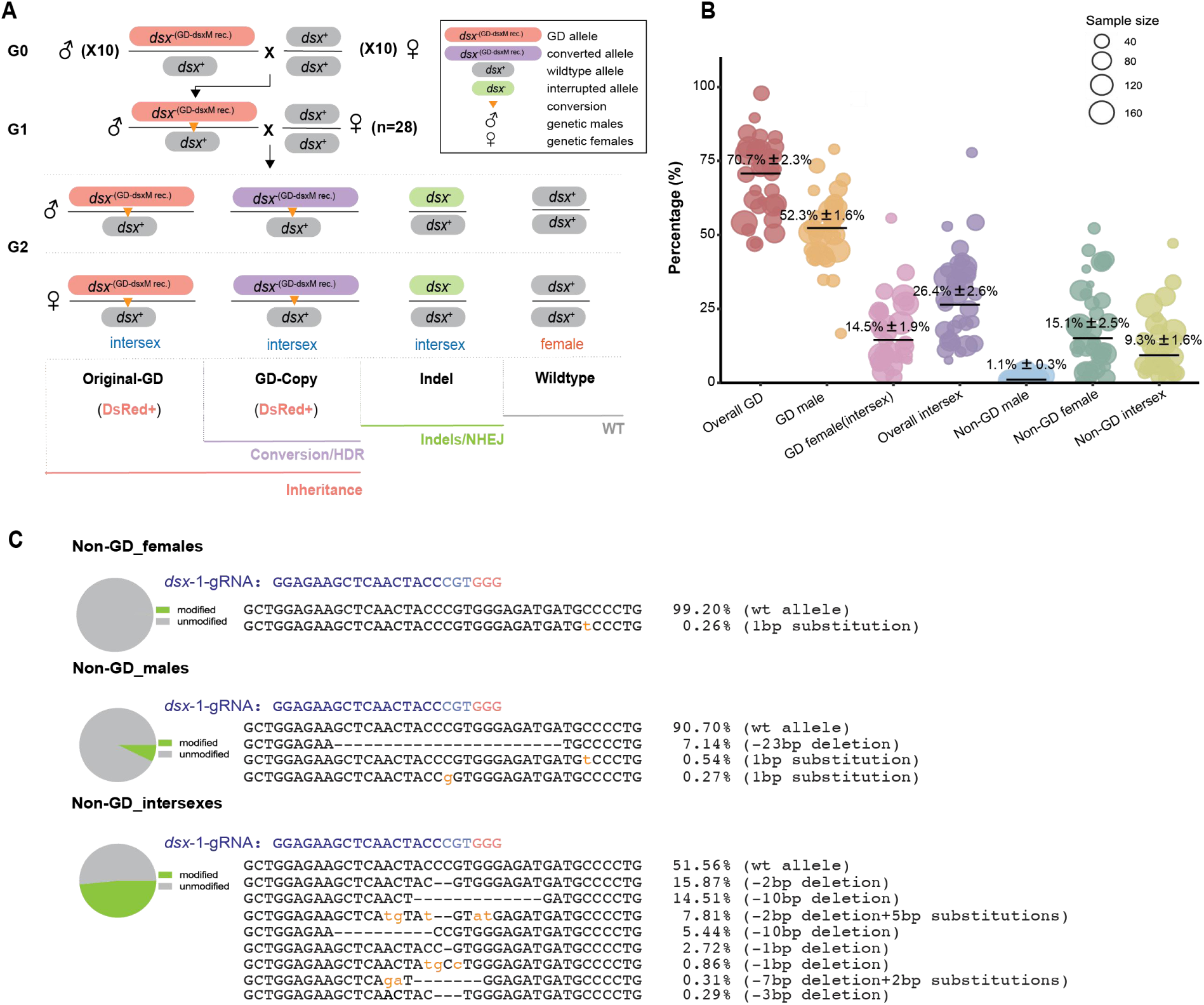
Inheritance and editing outcomes of the suppression gene drive targeting the *dsx* gene in *Cx. quinquefasciatus*. A: Schematic of the genetic crossing strategy used to assess gene drive efficiency. Transgenic males carrying the drive construct were crossed with virgin wild-type females to generate G1 heterozygous males, which were then single-pair crossed with virgin wild-type females to produce G2 progeny. Expected G2 phenotypic categories are illustrated below the crossing scheme. The *dsx* allele carrying the gene drive is shown in red, homing-converted alleles in purple, and resistance alleles in green. Orange inverted triangles represent potential homing events. **B:** Bubble chart showing the percentage of overall gene drive inheritance, GD-males, GD-females (or GD-intersexes), (calculated as the DsRed fluorescent individuals divided by total scored progeny), the percentage of overall intersexes (the total number of GD-intersexes and non-GD-intersexes), non-GD males, non-GD females and non-GD intersexes. A total of 33 independent germlines were analyzed. Mean values ± standard error of the mean (SEM) are indicated. Bubble size corresponds to the number of G2 progeny per cross. Raw phenotypic scoring data are provided in **Supplementary Data 3. C:** Editing outcomes at the *dsx*-gRNA-1 target site in non-GD individuals (males, females and intersexes) based on deep sequencing. Indel types and their relative frequencies are listed alongside.

Drive efficiency, measured as the proportion of DsRed-positive individuals among total G2 progeny, averaged 71%, exceeding the Mendelian expectation of 50% (**Fig. 2B**). The efficiency represents a notable improvement over our previously reported split gene drive, which achieved a maximum inheritance rate of 55%-60%^43^. Approximately 26% of G2 individuals exhibited intersex phenotypes, including GD intersexes resulting from ectopic *dsxM* expression of the drive construct in genetic females, and non-GD intersexes arising from end-joining (EJ)-induced mutations at *dsx* locus dominantly disrupted female development (**Fig. 2B**). Sanger sequencing confirmed that GD intersexes were heterozygous, carrying the drive on only one allele. On average, G2 progeny consisted of 52% GD males, 15% GD intersexes (formerly GD females with masculinized traits), ∼1% non-GD males, 15% non-GD females with wildtype alleles, and 9% non-GD intersexes harboring EJ-derived resistance alleles (**Fig. 2B**). These findings suggest that GD propagation was primarily driven by the males, while females carrying GD exhibited intersex phenotypes.

To further characterize editing outcomes, we pooled 30 non-GD females, males, and intersexes from G2 progeny (from multiple parents) for deep sequencing of the dual-gRNA target region. Unexpectedly, no editing was detected at the *dsx*-gRNA-2 site (driven by the U6:6 promoter), despite prior validation of U6:6 activity^18^ (**Supplementary Data 2**), suggesting that *dsx*-gRNA-2 was inactive. In contrast, variable editing was observed at the *dsx*-gRNA-1 site (**Fig. 2C**). Among non-GD intersexes, 48% of alleles were disrupted, consistent with heterozygosity at the *dsx* locus and supporting the hypothesis that indels at this site induce dominant sterility in females, likely by interfering with *dsxF* splicing. In non-GD males, 7% of alleles carried edits, with a 23-bp deletion being the most common indel. Normal male phenotypes were retained in these individuals, indicating a lack of dominant effect in males. No detectable edits were observed in non-GD females.

In GD-carrying individuals, edits at the *dsx*-gRNA-1 site were detected in 5.5% of GD males and 2.0% of GD intersexes (**Supplementary Fig. 1**), possibly due to somatic expression of Cas9 and gRNA. Although the molecular sex determination pathway in *Culex* remains poorly characterized, our deep sequencing results suggest that our target site at *dsx* may be essential for proper *dsxF* splicing. This implies that disruption of the both-sex exon can impair sex-specific splicing, leading to partial masculinization in females, perhaps due to shifting of splicing to the male isoform. In contrast, *dsxM* splicing appears unaffected by single-allele disruption, suggesting its regulation may be robust or independent of the edited site.

### Phenotypic and genotypic characterization of offspring

Phenotypic analysis of G2 individuals from single-pair cross revealed that all individuals inheriting the GD element developed as either males or intersexes. Genetic females carrying the *dsxM*-recoded drive allele exhibited masculinized features, including male-like plumose antennae, elongated maxillary palps, and malformed genitalia. Two distinct intersex phenotypes were observed: GD-intersex-I, characterized by laterally twisted claspers, and GD-intersex-II, with backward-rotated claspers (**Fig. 3A-c’ and d’**). In contrast, GD males were phenotypically indistinguishable from wild-type males. Among non-GD individuals, we observed wild-type males, wild-type females, and intersexes with varying degrees of masculinization. Wild-type individuals displayed clear sexual dimorphism (**Fig. 3A-a’ and b’**). Non-GD intersexes were classified into two categories based on external morphology. Non-GD-intersex-I individuals exhibited female-like traits, including pilose antennae, elongated maxillary palps, and cerci-like genitalia (**Fig. 3A-e’**). Non-GD-intersex-II individuals displayed masculinized features resembling those of GD-intersexes, including plumose antennae, abnormal maxillary palps, and malformed claspers (**Fig. 3A-f’**). Notably, non-GD-intersex-II individuals were rarely recovered, suggesting that this phenotype is relatively infrequent.

**Figure 3:**
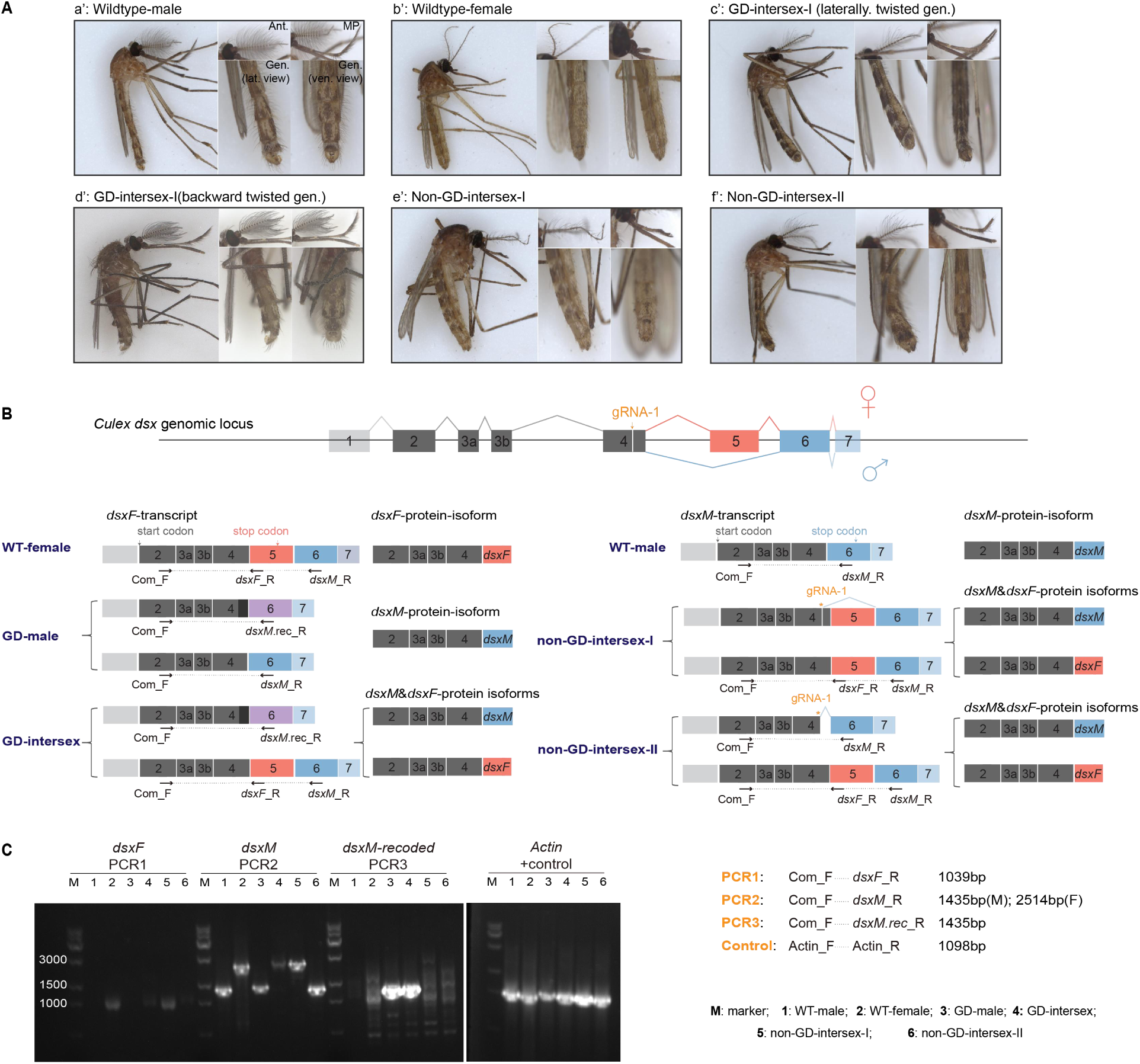
Phenotypic and genotypic characterization of offspring types. A: Sexual dimorphisms in antenna (Ant.), maxillary palps (MP), and genitalia (Gen.) structures across various offspring types. a’-wild-type male; b’-wild-type female; c’-GD intersex-I with laterally twisted genital claspers; d’-GD intersex-II with backward-rotated claspers; e’-non-GD intersex-I exhibiting partially feminized features; f’-non-GD intersex-II with masculinized features resembling those of GD intersexes. **B:** Schematic representation of *dsx* splice variants and corresponding protein isoforms detected in gene drive and non-drive offspring. Exons are not drawn to scale. **C**: Gel electrophoresis of RT-PCR products showing sex-specific *dsx* transcript expression across offspring types. Primer locations are indicated by dashed lines in panel B. *Culex Actin5C* gene (*CPIJ009808*) served as a positive control. Sample types and expected PCR product sizes are labeled next to the gel image.

Due to the absence of sex-specific genetic markers in *Cx. quinquefasciatus*, the genetic sex of non-GD intersexes could not be determined by standard PCR. To clarify the molecular basis of these phenotypes, we designed sex-specific primers targeting distinct *dsx* splice variants and performed RT-PCR using cDNA templates (**Fig. 3B**). GD males expressed both the endogenous *dsxM* transcript and the recoded *dsxM* transcript from the drive construct, confirming their heterozygosity (**Fig. 3C**). GD intersexes expressed both *dsxF* and recoded *dsxM* transcripts, indicating they were genetic females carrying the drive (**Fig. 3C**). Ectopic expression of *dsxM* likely contributed to their masculinized intersex phenotype. In contrast, non-GD intersexes expressed only endogenous splice variants. Non-GD-intersex-I individuals expressed *dsxF* but not *dsxM* (**Fig. 3C**), consistent with their more female-like appearance. Non-GD-intersex-II individuals expressed both *dsxF* and *dsxM*, correlating with their masculinized phenotypes that closely resembled those of GD intersexes (**Fig. 3C**). Sanger sequencing of non-GD-intersexes-I and non-GD-intersex-II revealed diverse indel variants at the *dsx*-gRNA-1 target site (**Supplementary Fig. 2**). Whether these indels are associated with the distinct phenotypes remains to be further investigated, particularly given the rarity of type-II individuals recovered and the limited understanding of the sex-determination cascade in *Culex* mosquitoes.

### Fitness cost evaluation of intersex mosquitoes

To assess the fitness costs associated with ectopic expression of the recoded *dsxM* element (in GD individuals) and *dsx* disruption via end-joining (in non-GD individuals), we evaluated the longevity, fertility, and hatchability of GD and non-GD intersexes derived from sibling offspring of GD males crossed with wild-type females to minimize batch effects. The following experimental groups were established: 1) 10 GD males X 10 wild-type females, 2) 10 GD intersexes X 10 wild-type males, 3) 10 non-GD intersex-I intersexes X 10 wild-type males (two biological replicates), and 4) a control group of 10 wild-type females X 10 wild-type males. Mating crosses for non-GD intersex-II individuals could not be established due to their low recovery frequency. However, given their phenotypic similarity to GD intersexes, their fitness was presumed to be similarly compromised. Survival was monitored daily, and blood feeding was performed between days 5 and 6 (**Supplementary Data 4**).

Survival analysis revealed that GD intersexes exhibited a reduced life span (14 days) compared to wild-type females (31 days) and wild-type males (26 days) (**Fig. 4A**). GD males showed a lifespan comparable to that of wild-type males (**Fig. 4A**). To assess their mating competitiveness, we introduced equal numbers of GD and wild-type males into mixed-sex cages containing virgin wild-type females. Since each *Culex* egg raft is typically laid by a single female, mating outcomes were evaluated based on the phenotype of offspring from individual rafts. Of the seven collected egg rafts, four produced DsRed-positive progeny, indicating successful mating by GD males (**Supplementary Data 5**). These results suggest that GD males remain viable and competitively mate with wild-type females. Interestingly, non-GD intersex-I individuals displayed extended lifespans (38 and 43 days in the two replicates) compared to wild-type females (31 days) (**Fig. 4A**). This could be related to their reduced rate of reproduction.

**Figure 4:**
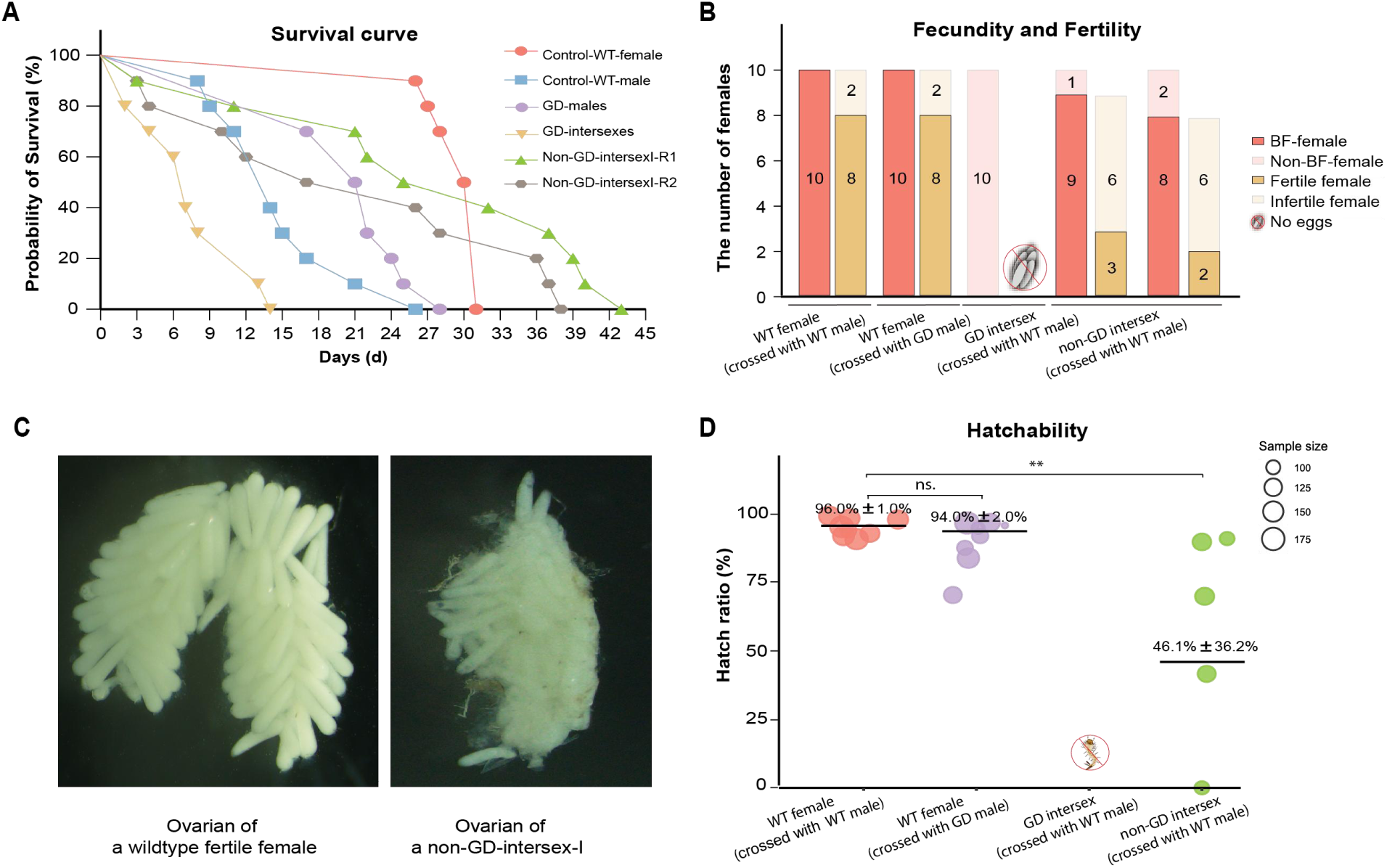
Fitness cost assessment of gene drive and non-gene drive individuals. A: Survival curves for wild-type males, wild-type females, GD-males, GD-intersexes and non-GD intersex-I groups. Two independent biological replicates were conducted for the non-GD intersex-I group. **B:** Fecundity and fertility measurements across different female types. Bars represent the number of blood-fed versus non-blood-fed females, and among those that are fed, the number of egg-laying (fertile) versus non-egg-laying (infertile) individuals, as determined by egg raft deposition. **C**: Representative images of ovarian structures in a fertile wild-type female and a non-GD intersex-I individual exhibiting oviposition failure. **D:** Hatch rates of egg rafts laid by different female types. Mean values ± standard error of the mean (SEM) are indicated. “ns” indicates no significant difference from the control group (WT female x WT male), and “**” indicates a significant difference (P<0.05). Raw data and statistical details are provided in **Supplementary Data 4**.

Fecundity and fertility were assessed by monitoring blood feeding and oviposition. All wild-type females crossed with either wild-type or GD-males successfully blood-fed, and 8 out of 10 laid eggs in both groups (**Fig. 4B**). In contrast, GD intersexes failed to blood-feed and did not lay eggs. Among non-GD-intersex-I individuals, 9/10 and 8/10 successfully blood-fed in the two replicates, respectively. However, only 3/9 and 2/8 blood-fed individuals laid eggs (**Fig. 4B**). A second blood-feeding attempt did not improve feeding rates, and some individuals died with undigested blood in their abdomens (**Supplementary Fig. 3A**).

Compared to the well-developed ovaries of fertile wild-type females, non-GD-intersex-I individuals with oviposition difficulties exhibited underdeveloped and morphologically abnormal ovaries (**Fig. 4C**). Dissection revealed retained eggs in the abdomen, some of which were undeveloped or darkened (**Supplementary Fig. 3B**). Hatchability assays showed that eggs from wild-type females mated with either wild-type or GD males had comparable hatch rates (**Fig. 4D**). In contrast, although non-GD-intersex-I individuals could lay a limited number of eggs, hatch rates were significantly reduced relative to wild-type females (**Fig. 4D**).

### Modeling performance of population suppression in *Cx. quinquefasciatus*

Our *Culex* suppression gene drive system can be classified as a RIDD-type strategy, as it targets the dominant *dsx* locus and relies on the release of transgenic males to propagate the drive and suppress the population by generating sterile intersex individuals. To evaluate the effectiveness of this system, we simulated releases of RIDD males at varying release ratios and monitored the number of fertile females (a key proxy for biting rates and disease transmission potential) on a weekly basis.

The release ratio refers to the weekly number of released mosquito males per generation (one generation = 3.17 weeks under normal circumstances), expressed as a fraction of the number of wild-type males present when the population is at carrying capacity. Modeling results show that although the RIDD system exhibits only moderate suppression strength, it remains effective in populations with low intrinsic growth rates (**Figure 5A**). This contrasts with previous models and experimental demonstrations^6,26^, largely due to the relatively modest total cut rate (drive conversion plus germline resistance formation) in our *Culex* drive, which leaves a substantial proportion of intact wild-type alleles. In addition, resistance alleles exhibit incomplete dominant sterility, further limiting suppression efficiency. Although RIDD males can eventually dominate the male population, the persistence of wild-type and partially fertile resistance alleles imposes a significant constraint on full population elimination^26^.

**Figure 5:**
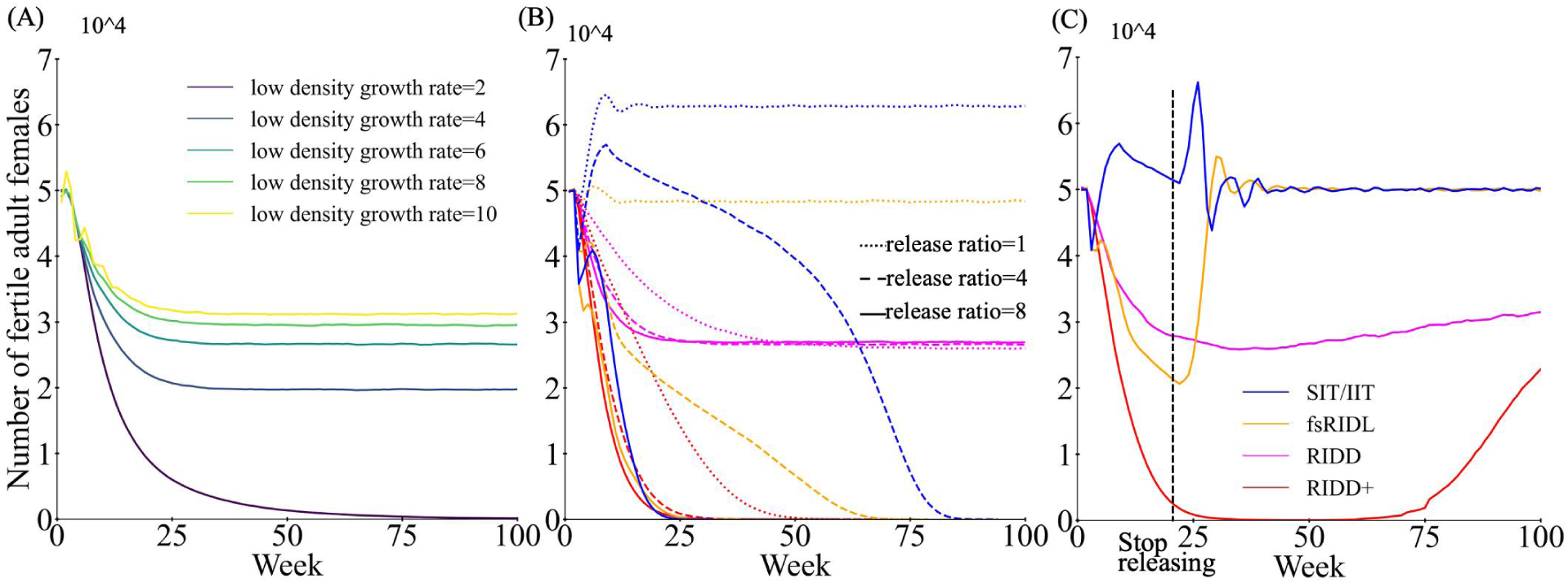
Modeling RIDD suppression gene drive in *Cx. quinquefasciatus*. Simulations were conducted by continuously releasing engineered males into a population containing 50,000 adult females at varying release ratios. The release ratio is defined as the number of engineered males per generation (3.17 weeks) relative to the size of the normal adult male population. **A:** RIDD males were released into populations with varying growth rates at low population density. **B:** Comparative performance of different genetic control strategies. RIDD+ is a hypothetical enhanced version of RIDD with an increased germline resistance rate (0.527 vs. 0.37). **C:** Suppression dynamics following a 20-week release of engineered males at a release ratio of Each data point represents the average of 10 replicates.

We next compared the performance of RIDD with other population suppression strategies, including the sterile insect technique (SIT) and female-specific RIDL (fs-RIDL)^50^. We also modeled a hypothetical RIDD+ drive with an increased germline cut rate (though still below the near-100% efficiency achieved in other species^43,51,52^) and complete dominant sterility conferred by resistance alleles. Our results indicate that under low release ratios, both RIDD and RIDD+ systems outperform SIT and fs-RIDL due to their greater persistence (**Figure 5B**). Notably, SIT is in particular vulnerable to overcompensation at low release ratios, which actually increases the population size. At higher release ratios, the advantage of the RIDD system diminishes, and the population still declines to nearly the same intermediate level. However, this limitation can be mitigated by increasing the level of germline cutting rates (as in RIDD+), enabling full population collapse even at low release ratios. Importantly, all these systems remain self-limited. When releases are conducted for only 20 weeks with a release ratio of four, both RIDD+ and RIDD systems suppress population rapidly, like fs-RIDL (**Figure 5C**). When releases cease, the population size returns to its original level more slowly compared to other control strategies.

## DISCUSSION

In this study, we established a population suppression gene drive system in *Cx. quinquefasciatus* by targeting a conserved region of the *dsx* gene and introducing a recoded *dsxM* sequence to restore male-specific isoform expression. This design ensures that GD-carrying females are converted into sterile intersexes, thereby enabling drive propagation through males while also contributing to the confinement of the system. Additionally, *dsx* disruption via end-joining in non-GD female offspring produced intersex individuals with substantial dominant fertility costs, indicating that *dsx*. These findings are consistent with observations in *D. melanogaster^6^* and *D. suzukii^34^*, where disruption of the female-specific exon leads to dominant female sterility, but contrasts with observations in *Anopheles* species, where disruption of *dsx* appears to be recessive when targeted at the female-specific exon or splice site^12,14^. This discrepancy may arise from differences in insect genera or from target site location: in *Culex*, disruption of a highly conserved region likely interferes with normal female-specific splicing, leading to a shift toward male-specific isoforms and masculinized intersex development. Importantly, the accumulation of resistance alleles in non-GD individuals, either in heterozygous or homozygous form, generates intersexes with substantially reduced fitness, accelerating population collapse.

Compared to our previously tested split gene drive system, which achieved a maximum inheritance rate of 60%^43^, the suppression drive reported here reached up to 71%, likely due to the use of a germline-specific *nanos* promoter, though site-specific and gRNA sequence-specific effects may also have contributed. Although a multiplexed gRNA strategy was employed, only one gRNA (*dsx*-1) proved functional. The inactivity of *dsx*-2 is unlikely due to the U6:6 promoter, which has demonstrated moderate editing activity in prior studies^18,43^. Instead, suboptimal target site selection is the probable cause. Notably, the homology arms were designed assuming both gRNAs would be functional. Since only *dsx*-gRNA-1 induced cleavage, the resulting mismatch between the homology arms and actual cut site may have reduced homing efficiency^38^. Further optimization of gRNA design would likely enhance drive performance. Even a simplified design using only *dsx*-gRNA-1, with homology arms precisely matching its cut site, should yield at least somewhat higher drive conversion rates.

The poorly characterized primary sex-determining signals and downstream regulatory pathways in *Culex* mosquitoes complicate genetic sex determination^42^. To address this, we used RT-PCR analysis of *dsx* splice variants to determine the genetic sex of intersex individuals and assess alternative splicing patterns. In GD females, the presence of the recoded *dsxM* element induced masculinized intersex phenotypes. In contrast, non-GD intersexes generated via end-joining formed two distinct phenotypic classes: Type I, with more female-like genitalia, and Type II, resembling GD intersexes with more male-like features. RT-PCR revealed that Type I intersexes expressed only the *dsxF* isoform, while Type II expressed both *dsxM* and *dsxF*. Sanger sequencing showed that these phenotypic differences may be associated with distinct indel types, but this remains inconclusive due to the limited sample size of Type II intersexes. It is hypothesized that disruption of the *dsxF* splice acceptor site may interfere with spliceosome recognition, leading to exon skipping and activation of the downstream male-specific exon. Although Type I intersexes retained partial fertility, their reduced fertility and hatchability still contributed to population suppression. This phenotype parallels findings from a suppression GD in *D. suzukii*, where hemizygous females exhibited normal mating behavior but markedly reduced fertility^34^.

Modeling results suggest that complete population suppression is not achievable with the current RIDD system unless the population has a very low intrinsic growth rate. However, success may be attainable with further modest improvements. The RIDD system performs with high effectiveness at low release ratios or when releases are limited to a short period, highlighting its potential to achieve substantial suppression even with a limited number of engineered individuals. To enhance efficacy at higher release ratios, improved Cas9 or gRNA expression is needed to approach near 100% cutting and ensure dominant sterility of resistance alleles.

A practical consideration is that producing RIDD males requires an additional outcrossing step to generate appropriately configured release individuals, which may introduce extra costs compared to strategies like fs-RIDL, where engineered males can be produced more directly^26^. Nonetheless, the outcrossing step also facilitates introgression of desired genetic backgrounds in the released males, an advantage shared with other homing type drives that can be more easily adapted to different genetic backgrounds.

Of note, the linkage between the drive and the male-determining locus contributes strongly to its performance. It allows greater persistence of the drive (and thus lower release ratios to achieve maximum effect) because lack of recombination ensures that the original drive allele is predominantly inherited by male progeny, sustaining the drive across generations. However, it also limits the overall power of the drive, as more females inherit wild-type alleles and resistance alleles that allow partial fertility. If germline cutting rates can be increased while maintaining low levels of resistance allele formation, most female progeny would become sterile, thereby preserving the drive’s benefits. Under these conditions, the system would function similarly to Y-linked editors targeting dominant genes, achieving effective population elimination even when drive conversion is low and enabling successful population elimination with very low release sizes^26,53^.

In summary, we have developed a self-limiting suppression gene drive targeting the *dsx* locus in *Culex* mosquitoes. Despite modest drive efficiency, modeling indicates that the system, with only modest improvements, can achieve effective suppression at low release ratios within a limited timeframe. This work expands the genetic toolkit for *Culex* vector control and lays the foundation for precise, sustainable, and ecologically responsible strategies to manage mosquito populations. Further optimization of gRNA activity and homing efficiency will be critical for maximizing the system’s effectiveness.

## METHODS

### *Culex* Mosquito Rearing and Maintenance

The *Culex quinquefasciatus* (Beijing strain, China) strain was kindly provided by Dr. Jianying Liu at Shenzhen Bay Laboratory. Mosquitoes were rearing at 27 ± 1°C, 75% relative humidity, under a 12-hour light/dark cycle in a Biosafety Level 2 (BSL-2) insectary room. Adults were maintained on a 10% sucrose solution. After mating, females were blood-fed on mice, and egg rafts were collected four days post-blood feeding. Larvae were fed with fish food pellets (Aquafin, China). Fluorescence and phenotypic assessments were performed using SZx2-ZB16 fluorescent microscope (Olympus, Tokyo, Japan). All procedures were carried out in accordance with protocols approved by the Institutional Biosafety Committee of Shenzhen Bay Laboratory and followed relevant ethical guidelines for animal research. Wastewater and used containers were frozen at −20°C for 48 hours before disposal as biohazardous waste.

### Plasmid Construction

All plasmids used in this study were generated using standard molecular biology techniques. Genomic DNA was extracted from approximately 10 adult wild-type *Cx. quinquefasciatus* mosquitoes using the Animal Tissue DNA Extraction kit (Vazyme, #DC102-01, China). The *dsx* gene was amplified and TA-cloned to serve as the plasmid backbone. Two *dsx*-targeting gRNA fragments (*dsx*-gRNA-1 and *dsx*-gRNA-2) were synthesized by annealing complementary oligonucleotides and inserted into the double-BbsI sites of the *Cq-*U6_1_2XBbsI-gRNA plasmid (Addgene #169238) and *Cq-*U6_6_2XBbsI-gRNA plasmid (Addgene #169323) plasmids, respectively. The gRNA scaffold was modified by incorporating a 5-bp loop structure and a T-C polymorphism to enhance transgenesis, as described in our previous studies^18^. The *Cq-* U6_1_*dsx*-gRNA-1 and *Cq-*U6_6_*dsx*-gRNA-2 cassettes were amplified and cloned into the *dsx* homology arm backbones flanking the gRNA cut sites. The *nanos*-Cas9 component was amplified from the *Cq*-*nanos*-Cas9 plasmid (Addgene #169348). Plasmids were assembled using Gibson Assembly (NEBuilder HiFi DNA Assembly Master Mix, New England Biolabs, # 2621). Following transformation into *E. coli DH5a* chemically competent cells (Sangon Biotech, #B528413, China), positive clones were verified by restriction enzyme digestion and Sanger sequencing. A list of primers used for plasmid construction is provided in Supplementary Table 1. Plasmid sequence is available in GenBank, with accession numbers listed in the Data Availability section.

### Fitness Cost Evaluation

Various mating crosses were established among different offspring categories. Survival, blood-feeding success, oviposition, and egg hatch rates were monitored daily. For non-GD intersex individuals with oviposition difficulties, abdominal dissections and imaging were performed using a microscope (SZMN, China) to assess reproductive anatomy and detect retained or abnormal eggs.

### Genotyping and RT-PCR

Genomic DNA (gDNA) was extracted from mosquito legs using STE squishing buffer (Solarbio, China) supplemented with proteinase K (50:1 ratio), followed by a PCR-based digestion cycle (65°C for 45 min, 95°C for 15 min, and held at 16°C). The extracted gDNA was used as a template for PCR amplification with primers designed to cover *dsx*-gRNA-1 and *dsx*-gRNA-2 target regions. PCR products were analyzed by Sanger sequencing to detect end-joining induced resistance alleles, and indel variants were quantified using the ICE CRISPR analysis tool^54^. For RT-PCR analysis, total RNA was extracted from adult mosquitoes with distinct phenotypes using the phenol-chloroform method^55^. First strand cDNA synthesis was performed using commercial cDNA synthesis kit (Vazyme, #R312-02, China). RT-PCR was conducted with gene-specific primers, and amplification products were visualized by agarose gel electrophoresis. All primers used are listed in Supplementary Table 1.

### Amplicon sequencing

Approximately 30 individuals from each G2 offspring category with distinct phenotypes were collected for analysis. Genomic DNA was extracted using the Animal Tissue DNA extraction Kit (Vazyme, #DC102-01, China). PCR amplification targeting the gRNA editing sites was performed, and products were purified using a gel DNA extraction kit (Vazyme, #DC301-01, China) in preparation for deep sequencing (conducted by Qingke Biotechnology). Sequencing data were analyzed using CRISPResso2 pipeline^56^. Primer sequences used for amplicon sequencing are listed in Supplementary Table 1.

### Statistics and Graphical Representation

All figures were generated using GraphPad Prism 9 and Adobe Illustrator. Mosquito images were captured using a VHX-7000C imaging system (KEYENCE). Inheritance ratio and hatch ratio analyses were performed in R using a “batch effect” approach, employing a generalized linear mixed model fitted by maximum likelihood to account for variance between cross batches^12^. No statistical method was used to predetermine sample size, and no data were excluded from the analyses. Hatchability comparisons among different groups were conducted using one-way ANOVA with Tukey’s multiple comparison test.

### Modelling of the Culex quinquefasciatus RIDD

Since most mosquito vectors share broadly similar life cycles, the effectiveness of releasing RIDD mosquitoes into wild populations was evaluated using a previously developed mosquito-specific model with weekly time steps^57,58^ implemented in the forward genetic simulation software SLiM 4^59^. Female mosquitoes typically mate only once and store sperm for subsequent egg production. In this model, we incorporated a 5% probability per week that a female may remate, allowing for the possibility of additional mating beyond the initial one. Females mate randomly with any fertile male in the population. Each week, each mated female has a 50% possibility of producing offspring. The number of eggs laid by a mated female follows a Poisson distribution with an average of 50.

In our model, the first two weeks for an individual represent the juvenile stage of mosquito development. During this period, the newly hatched larvae experience density-dependent competition, which is modeled according to the following equations:

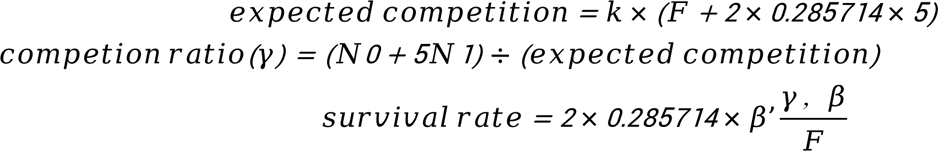

where *K* is the number of adult females in the population at the carrying capacity, *F* is the expected number of offspring produced by each adult female, N_0_ and N_1_ represent the number of new larva (age 0) and older larva (age 1) separately, *β*’(*r*,*β*) represents a linearly decreasing function of *r*, where *β* denotes the intrinsic growth rate under low-density conditions. Adult mosquitoes in the model are subject to an age-dependent survival rate. For adult males, the weekly survival probabilities are [2/3,1/2,0] indicating a maximum lifespan of three weeks. For adult females, the survival probabilities are [5/6, 4/5, 3/4, 2/3, 1/2, 0], allowing them to live for up to six weeks.

We implemented multiple releases of drive-carrying male heterozygotes for the RIDD system, in which ideally, both nonfunctional resistance alleles and drive allele exhibit dominant female sterility^6^. However, to match experimental observations, females with nonfunctional resistance alleles can lay eggs at a rate of 31%, and if they do lay eggs, they will average 62% as many as a wild-type mosquito. We also modeled a variant that assumed dominant sterility of nonfunctional resistance alleles. For comparison, we also simulated population suppression strategies based on the Sterile Insect Technique (SIT) and the release of males homozygous for a female-specific RIDL (fs-RIDL) construct. In the SIT strategy, females that mate with released sterile males produce no viable offspring. In the fs-RIDL strategy, the construct carries a dominant lethal gene that causes female offspring to be nonviable. This occurs at the early larva stage before competition takes place (based on an existing construct^50^). Not fitness costs were modeled, other than intended costs by the drive.

Simulations were conducted for up to 317 weeks (approximately 100 generations) unless the number of fertile females reached zero earlier. To establish a baseline equilibrium, wild-type mosquitoes were allowed to reproduce randomly for 10 weeks prior to the introduction of drive-carrying mosquitoes. Given that functional resistance can potentially be avoided through the use of multiple gRNAs or by targeting a highly conserved site^14,37,38^, our model included only nonfunctional resistance alleles (r2).

Simulations were conducted using the High-Performance Computing Platform at the Center for Life Sciences, Peking University. Data processing and figure generation were performed using Python. Default parameters for all models are listed in Table 1, and the corresponding SLiM scripts and raw data are available on Github (https://github.com/jchamper/Culex-RIDD).

**Table 1.** Default parameters for the model.

| Drive system | Parameter | Default value |
| --- | --- | --- |
| RIDD only | Drive conversion rate | 0.414 |
|  | Germline resistance formation rate | 0.37 |
|  | Recombination rate | 0.01876195 |
|  | Female fertility with resistance | 0.31*0.62 |
| RIDD+ | Germline resistance formation rate | 0.9 |
|  | Female fertility with resistance | 0 |
| All models | Low density growth rate | 6 |
|  | Number of adult females | 50000 |

## DATA AVAILABILITY STATEMENT

The plasmid sequence of the suppression gene drive construct in this manuscript is deposited into the GenBank database and available from the authors upon request. GenBank accession numbers for the deposited plasmid is: P69_Cq_suppression GD (NCBI PV864762). Code used for modeling is available on GitHub (https://github.com/jchamper/Culex-RIDD).

## ACKNOWLEDGEMENTS

The research reported in this manuscript was supported by the Zhejiang University, Key Laboratory of Biology of Crop Pathogen and Insects of Zhejiang Province, College of Agricultural and Biotechnology, China, by Shenzhen Bay laboratory, Shenzhen, China, by National Natural Science Foundation of China grants number 82202559 (to XF), 82372289 (to FL) and 32270672 (to JC), by Zhejiang Province Science Foundation MS25C140016 (to XF), and by the Center for Life Sciences (to JC).

## AUTHOR CONTRIBUTIONS

XF and FL conceived the project; XF and VMG. contributed to the design of the experiments; XF and JD performed the experiments and contributed to the data collection and analysis; YL performed the modelling and analysis; XF, FL, JD, YL, XC and JC drafted the manuscript. All authors revised the manuscript.

