## Supplementary information for "Self-limiting population suppression gene drive in the West Nile vector mosquito, *Culex quinquefasciatus*"

[1] Key Laboratory of Biology of Crop Pathogen and Insects of Zhejiang Province, College of Agricultural and Biotechnology, Zhejiang University, Hangzhou, China
[2] Institute of Infectious Diseases, Shenzhen Bay Laboratory, Shenzhen, China
[3] Center for Bioinformatics, School of Life Sciences, Center for Life Sciences, Peking University, Beijing, China
[4] Center for Infectious Diseases, Department of Epidemiology, Human Genetics, and Environmental Sciences, School of Public Health, University of Texas Health Science Center, Houston, TX, USA
[5] Section of Cell and Developmental Biology, University of California San Diego, La Jolla, CA, USA
[6] Guangdong Provincial Key Laboratory of Infection Immunity and Inflammation, Shenzhen, China
[#] Authors contributed equally

**Keywords**: *Culex* mosquitoes, population suppression, gene drives, *doublesex*, RIDD.

### Supplementary Figures

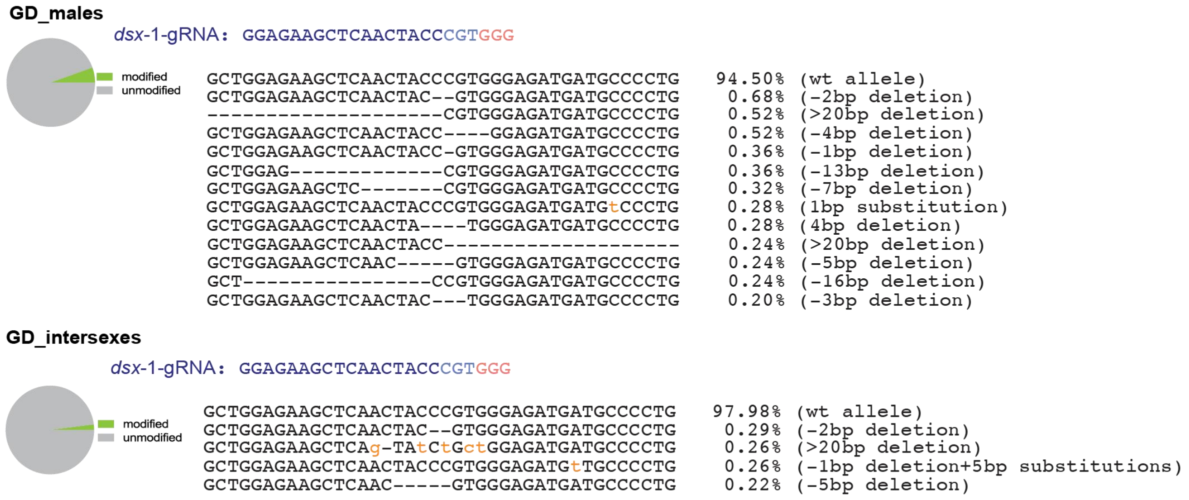


**Supplementary Figure 1: Editing outcomes at *dsx*-gRNA-1 target site in GD individuals**. Deep sequencing analysis (Supplementary Data 3) of GD males and intersexes revealed editing outcomes at the *dsx*-gRNA-1 target site. The proportions of modified versus unmodified alleles are shown as pie charts (left), while the types and frequencies of observed indels are listed on the right.

```
Wild-type
                                                  dsx-1-gRNA
TACATCAGGGGCATCATCTCCCACGGGTAGTTGAGCTTCTCCAGCAGATACTGAGATCGTTTCACCAGTTCATCG      (Wild-type allele)

Non-GD-intersex-type I (non wild-type alleles)

TACATCAGGGGCATCATCTCCCAC--GTAGTTGAGCTTCTCCAGCAGATACTGAGATCGTTTCACCAGTTCATCG      (-2bp deletion)
TACATCAGGGGCATCATCTCC-------tGaTGAGCTTCTCCAGCAGATACTGAGATCGTTTCACCAGTTCATCG      (-7bp deletion and 2bp substitution)
TACATCAGGGGCATC-------------AGTTGAGCTTCTCCAGCAGATACTGAGATCGTTTCACCAGTTCATCG      (-10bp deletion)
TACATCAGGGGCATCATCTC-------------------TCCAGCAGATACTGAGATCGTTTCACCAGTTCATCG      (-19bp deletion)
TACATCAGGGGCATCATCTCCCACG----------------------ATACTGAGATCGTTTCACCAGTTCATCG      (-22bp deletion)

Non-GD-intersex-type II (non wild-type alleles)

TACATCAGGGGCATCATCTCCCACGttgaGtTgGTTGAGCTTCTCCAGCAGATACTGAGATCGTTTCACCAGTTCATCG  (+4bp insertion and 2bp substitution)
```

**Supplementary Figure 2: ICE CRISPR analysis of *dsx*-gNRA-1 editing in non-GD intersex individuals**. Sanger sequencing traces from ten non-GD-intersex type I and one non-GD-intersex type II individuals were analyzed using ICE CRISPR. The identified non wild-type indel variants are listed.

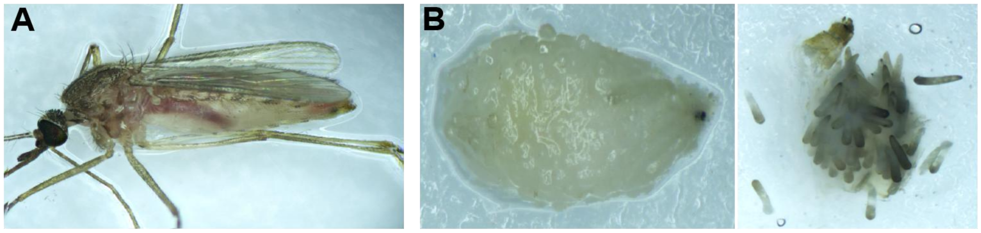


Dead non-GD intersex-I with undigested blood meal

Dissected ovarian structures showing undeveloped or darkened eggs in the abodomen of Non-GD intersex-I

**Supplementary Figure 3: Dissection images of non-GD intersex-I individuals with oviposition defects**. **A**: A dead non-GD intersex-I after blood feeding, showing retention of an undigested blood meal in the abdomen. **B:** Dissected abdomen revealing underdeveloped or darkened eggs, indicating impaired oogenesis or oviposition failure.

**Supplementary Tables**

**Supplementary Table 1-Primer List**

| **Name** | **Primer** |
|---|---|
| **Primers used for building a population suppression GD construct** | |
| 0493 | AAGTGCAACAACTCTCGGCTCGACCAAG |
| 0494 | ACAGTGGAGTAACGAAAAATGGCAGAG |
| 0526 | CATGAGTGGCATCATTTCCCAtGGaTAGTTGAGCTTCTCCAGCAGATACTGAG |
| 0582 | TGTACGTGATACTGAAGGGTGCCGACG |
| 0525 | CTCAGTATCTGCTGGAGAAGCTCAACTAtCCaTGGGAAATGATGCCACTCATG |
| 0527 | CTTGCAAGTCGTCGTCGTCGTCTGGTTGTACAATTAATCTAGTTGCATGCG |
| 0522 | GTTTTAGAGCTAtgctgGAAAcagcaTAG |
| 0523 | ACGGGTAGTTGAGCTTCTCCTCGGAGGGTGCCCTCGTTGC |
| 0528 | CGCATGCAACTAGATTAATTGTACAACCAGACGACGACGACGACTTGCAAG |
| 0529 | caaactcatcaatgtatcttaaagcttAAGGAACACCTGTTCCCTCTGCGAG |
| 0530 | CTCGCAGAGGGAACAGGTGTTCCTTaagctttaagatacattgatgagtttg |
| 0531 | ACCTTATTATCTTCGTAGATTTTGAcatactcggtggcctccccaccacc |
| 0522 | GTTTTAGAGCTAtgctgGAAAcagcaTAG |
| 0524 | TGATGTACGTGATACTGAAGTGTAGGGCCGGGAAAGTGCTTATATAG |
| 0532 | ggtggtggggaggccaccgagtatgTCAAAATCTACGAAGATAATAAGGT |
| 0583 | CGTCGGCACCCTTCAGTATCACGTACAAAAAAAAAAGCACCGACTCGGTGCCAC |
| Primers used for PCR-based verification of correct transgenesis | |
| X155 | CGCGAGTCAATATATCATTCGTG |
| X301 | ATGGTGCGCTCCTCCAAGAAC |
| X309 | TGAACTCCTTGATGACGTTCTTGGAGGAGCG |
| X310 | GGTTCGCCGAAGAAGAAGCGCAAGGTGTAAAATTG |
| X304 | ACAGTATaAAAAGTGCGTTAAAAAAACAAG |
| X51 | ATGGATAAGAAGTACTCGATCGG |
| X314 | TGGAAGAACGAATCATCCACCTTGGCCATC |
| X312 | AAAAATTTCACACCGTTCACGCAAGGCACTG |
| X313 | ACGTTATGTTCCTCCCAGTTCTGGCTGG |
| X157 | TGACCCAGGTCATGATCAACGACGACC |

| | |
|---|---|
| Primers used for *dsx-1* and *dsx*-2 Deep sequencing analysis | |
| X01 | GCTCAAAATAAACCTGATACGGGTG |
| X237 | GATCCAACCGCCACCGTTCCGTACC |
| Primers used for RT-PCR amplification of *dsxM*, *dsxF* and *dsxM*-recoded transcript fragments | |
| Com_F | X281-ATGGTTTCGCAAGATACCTGGATGG |
| *dsxF*_R | X315-CGTTTGTTTGCTCTCGGCAATAAAC |
| *dsxM*_R | X283-TCAGCGGATGGGGTGGATCACTACC |
| *dsxM*.rec_R | X298-GAGTGGATGTGGTGGGTCCGAACCGC |
| *Actin5C*-F | GTCGACAATGGATCCGGTATGTGCAAG |
| *Actin5C*-R | GAAGCACTTTCGGTGGACGATGGAC |

**Supplementary Data**

**Supplementary Data 1- Microinjection and transgenesis data for the population suppression gene drive**

This table represents raw counting data based on the presence of the DsRed fluorescent marker, indicating successful transgenesis. All hatched G0 individuals were sexed and pooled by sex, then crossed with wild-type individuals of the opposite sex. Egg rafts from each pool were hatched collectively, and the resulting offspring were screened and counted in pooled batches.

**Supplementary Data 2- Deep sequencing analysis**

Summary of the CRISPRessp2 batch analysis of allele editing frequencies based on deep sequencing data from categorized samples collected in GD experiments. The calculated allele editing frequencies are presented in **Fig. 2C** and **Supplementary Fig 1**. The data are organized into the following file tabs:

1. Allele type and frequency at *dsx*-gRNA-1 editing site
2. Allele type and frequency at *dsx*-gRNA-2 editing site

**Supplementary Data 3- Suppression gene drive data**

This dataset contains raw counts of G2 progeny with phenotypic scoring for DsRed+ males, DsRed+ intersexes, non-fluorescent males, females and intersexes. Inheritance of the gene drive element was tracked using DsRed fluorescence. Two independent batches of gene drive experiments were conducted at different times, and the resulting data were pooled for analysis. Transgene inheritance rates were calculated for each single-pair cross with mean values and standard deviations summarized in the table. Data were analyzed using a “batch effect” approach in R, employing a generalized linear mixed model fitted by maximum likelihood to account for variance between cross batches. These results are presented in **Fig 2B**.

**Supplementary Data 4- Raw data on the fitness costs of gene drive and non-gene drive intersexes**

This dataset includes raw measurements of adult survival, female blood-feeding behavior, egg laying performance, and hatch rates per egg raft from various genetic crosses. The summarized results are shown in Fig. 4A-C. Each tab corresponds to a specific cross scheme as follows:

1. **Control group**- 10 wild-type males X 10 wild-type female
2. **GD♂ × WT♀**- 10 gene drive (GD) males X 10 wild-type females
3. **GD intersexes × WT♂**-10 GD intersexes X 10 wild-type males
4. **Non-GD intersexes × WT♂**- 10 non-GD intersexes X 10 wild-type males (two biological replicates).

5. Statistical analysis details

**Supplementary Data 5-Mating competition between gene drive (GD) males and wild-type males**

This table presents the results of mating competition assays between GD and wild-type males. Ten newly emerged GD males and 10 wild-type males were pooled together with 10 virgin wild-type females. After 3 days of cohabitation, females were blood fed, and oviposition was induced 3 days post-feeding. Egg rafts were collected and hatched individually. Fourth instar larvae were screened under a fluorescence microscope, and the presence of DsRed+ marker was recorded to evaluate successful mating.

**Supplementary method**

**Cloning details for the plasmid constructs**

**P69_*Culex.q* population suppression gene drive (GD) construct**

The homology arms were amplified from *Culex quinquefasciatus doublesex* gene using genomic DNA and cloned into a TOPO vector to serve as the backbone for the GD construct. The recoded *dsxM* fragment was synthesized by GenScript, incorporating an *Anopheles gambiae Kh* 3'UTR. The *nanos*-Cas9 fragment was amplified from the plasmid pVMG0212 (Addgene #169348). The *U6:1_dsx*-gRNA-1 fragment was amplified from the Cq_*U6:1_dsx*-gRNA-1-LOOP plasmid modified from pVMG0302 (Addgene #169369). The *U6:6_dsx*-gRNA-2 fragment was amplified from the *Cq_U6:6_dsx*-gRNA-2-LOOP plasmid modified from pVMG0303 (Addgene #169370). The *Opie2*-DsRed-SV40_3'UTR fragment was amplified from pVMG0252 (NCBI accession # MW925705) and assembled with other components using Gibson Assembly. All primers used in this construct are listed in Supplementary Table 1.

**Plasmid sequences and availability:**

The plasmid sequence of the *Culex quinquefasciatus* population suppression gene drive (GD) construct generated in this study has been deposited in the GenBank database with accession number PV864762.
